# Human hippocampal ripples signal encoding of episodic memories

**DOI:** 10.1101/2022.10.03.510672

**Authors:** John J. Sakon, David J. Halpern, Daniel R. Schonhaut, Michael J. Kahana

## Abstract

Recent human electrophysiology work has uncovered the presence of high frequency oscillatory events, termed ripples, during awake behavior. This prior work focuses on ripples in the medial temporal lobe (MTL) during memory retrieval. Few studies, however, investigate ripples during item encoding. Many studies have found neural activity during encoding that predicts later recall, termed subsequent memory effects (SMEs), but it is unclear if ripples during encoding also predict subsequent recall. Detecting ripples in 116 neurosurgical participants (n = 61 male) performing an episodic memory task, we find insignificant ripple SMEs in any MTL region, even as these regions exhibit robust high frequency activity (HFA) SMEs. Instead, hippocampal ripples increase during encoding of items leading to recall of temporally or semantically associated items, a phenomenon known as clustering. This subsequent clustering effect (SCE) arises specifically when hippocampal ripples occur during both encoding and retrieval, suggesting that ripples mediate the encoding and future reinstatement of episodic memories.

## Introduction

Decades of work in animal models have identified discrete, high frequency events in MTL, termed ripples (*1*). This animal work has suggested a specific role for hippocampal ripples in memory formation during learning and offline replay (*1*) and more recently memory retrieval (*2*). A series of recent studies have investigated ripples in human intracranial recordings (see Liu *et al.* (2022) for a review (*3*)). Many of these investigations relate medial temporal lobe (MTL) ripples and memory retrieval, with ripple rate increasing just before participants vocalize recalls (*4–11*). The few studies that have reported ripple rates during memory encoding, however, find conflicting evidence regarding their relation to subsequent recall. One study finds an increase in ripple rates for subsequently recalled items 0.7-1.5 s into item presentation (*7*), while the other finds ripple increases only after item presentation (*5*).

A separate but related body of research has shown that *≥*60 Hz spectral power (often termed high frequency activity [HFA] or fast-gamma oscillations) predicts subsequent recall (*12–15*). Indeed, numerous intracranial studies using HFA detectors distinguish subsequently recalled and not-recalled items, termed a subsequent memory effect (SME) (*7, 16, 17*). The overlapping frequency ranges used to detect HFA and ripples raise questions about whether and how these signals may be related to one another (*18*).

Recent human intracranial studies find hippocampal ripples preferentially occur during recall of episodic memories (*6, 8*). In particular, Sakon & Kahana (2022) demonstrate that hippocampal ripples signal reinstatement of context during memory retrieval, a mechanism considered crucial to the “jump back in time” phenomenology of episodic memory (*19*). Meanwhile, theories of ripple function suggest memory formation and memory retrieval share mechanisms, as neural activity reinstated during memory retrieval overlaps with activity repeatedly reinstated during consolidation (*2, 10*). Considering these two ideas, do ripples also reflect context reinstatement during the formation of episodic memories? For example, if you attend a Philadelphia Phillies game and enjoy a cheesesteak, future cheesesteak orders may retrieve the context of the event, which will lead to reinstatement of memories clustered with the game (*20*). We hypothesize that during the formation of the memory, hippocampal ripples signal engagement of episodic memory mechanisms, which strengthen the association between the item (cheesesteak) and context (Phillies game). This association subsequently increases the likelihood of reinstating the context of the game when later cued by cheesesteak. Previous work has shown evidence of this phenomenon, termed a subsequent clustering effect (SCE), using HFA in hippocampus (*21*), hinting that ripples may underlie subsequent clustering.

Analyzing intracranial EEG recordings of 116 participants (233 sessions) performing a delayed free recall task, we ask if ripples show an SME or SCE in the hippocampus and surrounding cortical regions in MTL. We partitioned our data into two parts: an initial *∼*35% of participants for developing initial hypotheses and analyses, and a second part held out so that we could confirm our findings with the whole dataset. We pre-registered the initial hypotheses and figures supporting them on the Open Science Framework (OSF; https://osf.io/e98qp). Therefore, we defined the analysis parameters for our figures in the pre-registration based on the first part of the dataset, and here we present figures and statistics on the full dataset as well as statistics for the held out data for the key hypotheses (*Methods*). A full breakdown of the outcomes for these initial hypotheses has also been registered on the OSF (https://osf.io/aue9c). The analyses build the case that awake, hippocampal ripples specifically signal the formation of episodic memories. First, we do not find a significant ripple SME throughout any MTL regions despite replicating an SME for HFA. However, when partitioning words into those that lead to subsequent clustering of recalls vs. those that do not, we find a significant ripple subsequent clustering effect (SCE) specifically in the hippocampus. Evidencing its role in task performance, participants with a stronger hippocampal ripple SCE exhibit increased clustering and superior memory. Finally, we show that hippocampal ripples during memory formation lead to subsequent clustering precisely when ripples also occur prior to word recall, implying the SCE incites ripple-mediated reinstatement.

## Materials and Methods

### Human participants

Comprising the dataset are intracranial recordings from 116 adult participants (n=61 male) in the hospital for drug-resistant epilepsy surgery with subdural electrodes placed on the cortical surface or within the brain to localize epileptic activity. Collaborating hospitals include Thomas Jefferson University Hospital (Philadelphia, PA), University of Texas Southwestern Medical Center (Dallas, TX), Emory University Hospital (Atlanta, GA), Dartmouth-Hitchcock Medical Center (Lebanon, NH), Hospital of the University of Pennsylvania (Philadelphia, PA), Mayo Clinic (Rochester, MN), and Columbia University Hospital (New York, NY). All participants consented to research under a protocol approved by the Institutional Review Board at the University of Pennsylvania via a reliance agreement with each hospital.

### Experimental design and Statistical Analysis

Participants were tested on a delayed free recall task in which each “list” comprised viewing a sequence of common nouns with the intention to commit them to memory. The task was run at bedside on a laptop and participants were tasked to finish up to 25 lists for a whole session or 12 lists for a half-session. The free recall task consisted of four phases per list: countdown, encoding, distractor, and retrieval (**Fig. 1a**). Each list began with a 10-second countdown period with numbers displayed from 10 to 1. For encoding, participants were sequentially presented 12 words centered on the screen that were selected at random–without replacement in each whole session or two consecutive half sessions–from a pool of 300 high frequency, intermediate-memorable English or Spanish nouns (http://memory.psych.upenn.edu/WordPools (*22*)). Each word was presented for 1.6 s with a jittered 0.75-1.2 s (randomly sampled uniform distribution) blank screen shown after each word. After encoding was a distractor period where participants performed 20 seconds of arithmetic math problems to disrupt their memory for recently-shown items. Math problems were of the form A+B+C=??, where each letter corresponds to a random integer and participants typed their responses into the laptop keyboard. The final phase is retrieval, in which participants had 30 seconds to recall as many words–in any order–from the most recent list as possible. Retrieval began with a series of asterisks accompanied by a 0.3 s, 60 Hz beep to signal for the participants to begin vocalizing recalled words. Vocalizations were recorded and later annotated offline using Penn TotalRecall (http://memory.psych.upenn.edu/TotalRecall) to determine correct and incorrect recalls. For each session the participant began with a practice list of the same words that we do not include in the analysis.

**Figure 1.**
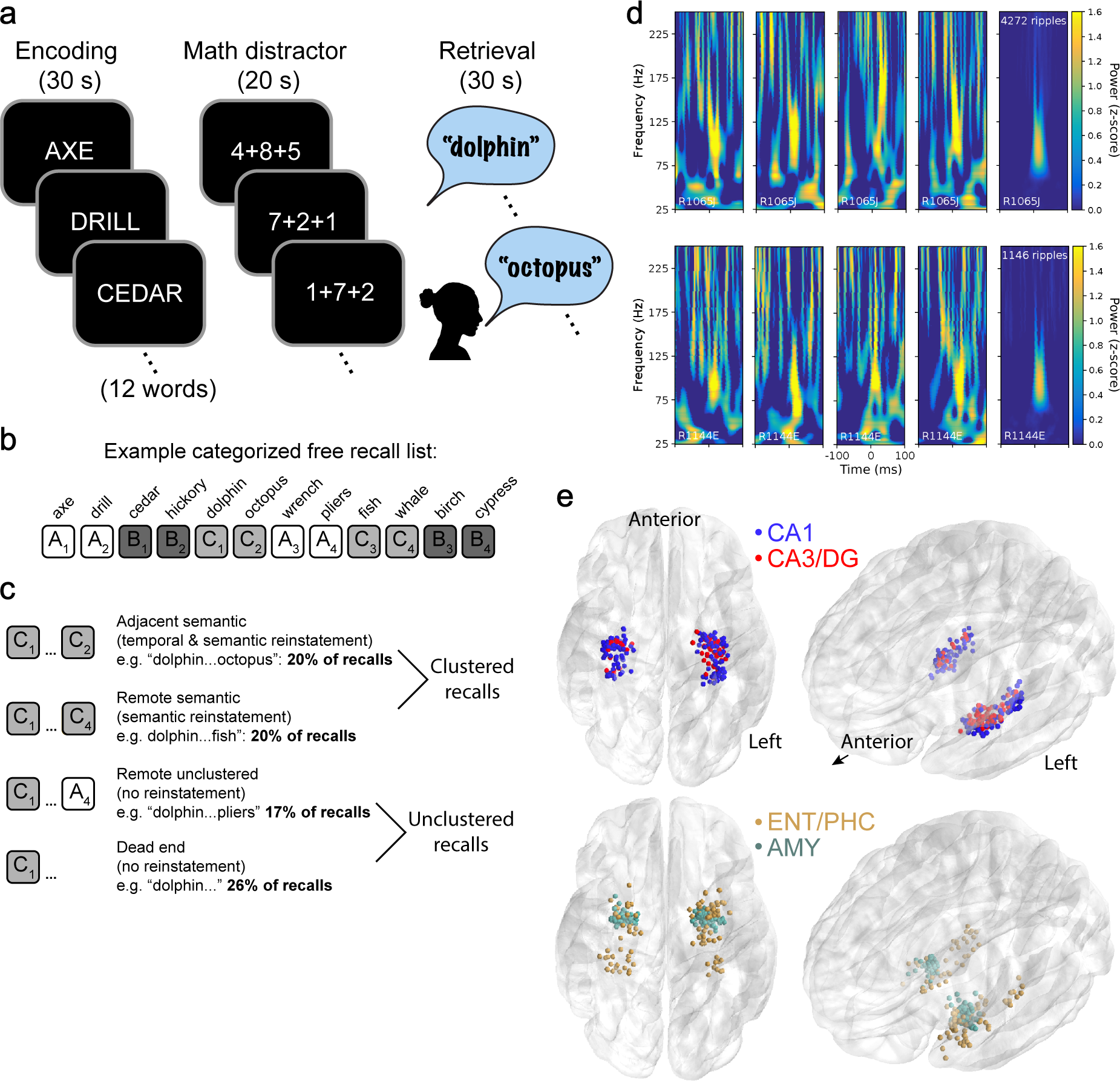
Free recall task design and ripple detection details. **(a)** Task diagram of delayed free recall, in which participants perform a math distractor in between word presentations and a retrieval period. **(b)** Structure of categorized word lists used in this task variant. A, B, and C are each semantic categories (tools, trees, and sea animals in this case). The two pairs of words from the same category are never shown back-to-back **(c)** Types of recall transitions in the categorized free recall task and percentage of recalls that lead to each. Note that adjacent, non-semantic transitions are only 3% of recalls due to the semantic nature of the task so are not analyzed. **(d)** Each row displays EEG spectrograms aligned to the start of ripples occurring during word presentation for two participants with hippocampal CA1 electrodes. The first four columns show single trial examples while the fifth column shows the average across all ripples during word presentation for all CA1 electrodes in all sessions for each participant. **(e)** Electrode bipolar pair midpoint localizations for all participants performing catFR. Shown are hippocampal subfields CA1 and CA3/dentate gyrus (CA3/DG), entorhinal (ENT) and parahippocampal (PHC) cortex, and amygdala (AMY).

All analyses in this manuscript are done on a variant called categorized free recall, in which each list is comprised of words with semantic relationships. For every whole session (or consecutive half sessions), words were drawn from a pool of 300 that included 12 words each from 25 categories created using Amazon Mechanical Turk to crowdsource typical exemplars for each category (*22*). For each list, three semantic categories were randomly chosen, and the four words from each category were presented sequentially in pairs. Pairs from the same category were never shown back-to-back (in other words, the four words from the same category were never shown in a row). This setup allowed us to study both adjacently (same pair) and remotely presented words from the same category.

A unique feature of intracranial data is that patients can vary dramatically in how much data they contribute to a particular cell in our statistical design. To equally weight patients who vary dramatically in the number of observations they contribute, we include both fixed and random effects in linear mixed effects models for all statistical tests to account for the varying effect sizes amongst patients. We present effects using *β* coefficients with standard errors and use a Wald test to evaluate statistical significance. Equations for each test are presented separately in the **Equations** section. We correct for multiple comparisons across brain regions using the Benjamini–Hochberg procedure for controlling false discovery rate (FDR), which is appropriate for positively correlated data such as brain activity during task performance.

### Intracranial electroencephalogram (iEEG) recordings

iEEG was recorded from macroelectrodes on subdural grids and strips (intercontact spacing 10.0 mm) or depth electrodes (intercontact spacing 3-6 mm) using DeltaMed XlTek (Natus), Grass Telefactor, Nihon-Kohden, Blackrock, or custom Medtronic EEG systems. Signals were sampled at 500, 512, 1000, 1024, 1600, 2000 or 2048 Hz and downsampled using a Fourier transformation to 500 Hz for all analyses. Initial recordings were referenced to a common contact, the scalp, or the mastoid process, but to eliminate possible system-wide artifacts and to better isolate localized high frequency signals we applied bipolar rereferencing between pairs of neighboring contacts. Bipolar referencing is ideal as the spatial scale of ripples is unlikely to exceed intercontact spacing of our recordings (3-10 mm) (*9*). Line removal is performed between 58-62 using a 4th order Butterworth filter (120 Hz is in our sensitive ripple range and we did not find artifacts in these frequencies).

### Ripple detection

Detection of ripples is identical to our previous work, where we performed numerous control analyses to ensure the detector is robust to vocalization artifacts, frequency window selection, correlations across channels, and seizurogenic activity (*4*), and is based on prior human work (*5, 23*). Briefly, local field potential from bipolar iEEG channels is bandpass Hamming filtered from 70-178 Hz, Hilbert-enveloped, squared, smoothed, and normalized to find candidate events exceeding 3 standard deviations (SD) that are expanded to find their duration above 2 SDs. Events are considered ripples if the expanded duration is between 20 and 200 ms and not within 30 ms of another expanded event (in which case the events are merged). To avoid pathological interictal epileptiform discharges (IEDs), LFP is bandpass Hamming filtered from 25-58 Hz rectified, squared, smoothed and normalized to detect events 4 SD above the mean. Ripples within 50 ms of an IED event are removed.

We treat ripples as discrete events (**Fig. 2a**) with the timestamp set to the beginning of each ripple (**Fig. 1d**). The average power of events is *∼*90 Hz, although individual events peak throughout the 70-178 Hz range (**Fig. 1d** shows 8 single ripple examples). Most participants had multiple MTL contacts within their montage, thereby providing iEEG recordings from multiple channels for every word presentation. As with previous work (*4, 5, 9*), since the spacing of clinical electrodes (3-10 mm) is much farther than ripples are expected to travel in the brain (*<*0.2mm, (*24*)), we consider each presented word for each channel as a separate “trial”. To ensure ripples are not double-counted across neighboring channels we use a combination of automated channel and session removal (by measuring correlations across trials and channels, respectively) and manual inspection of raster plots (**Fig. 2a**) as detailed in previous work (*4*).

**Figure 2.**
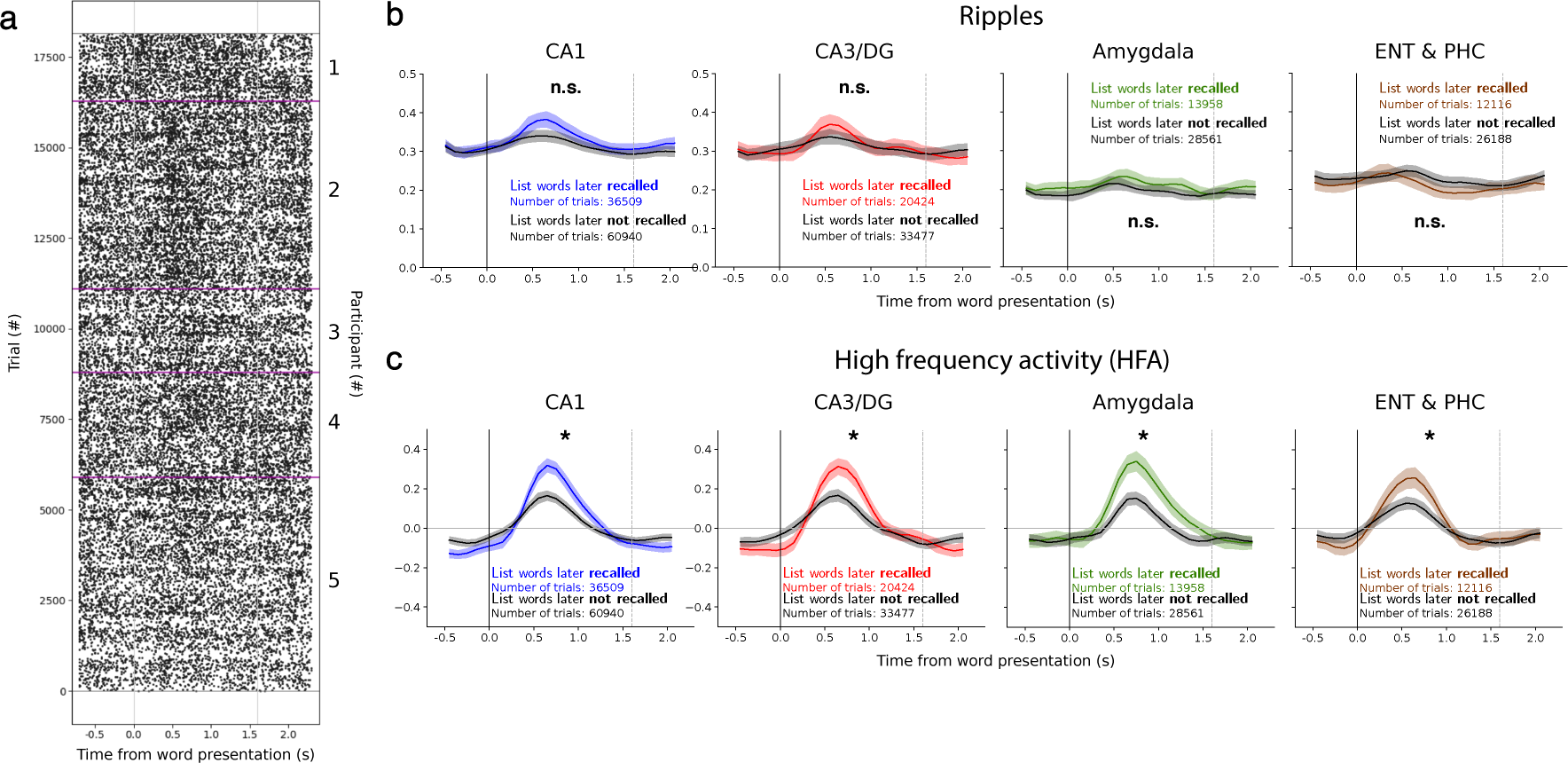
High frequency activity (but not ripples) shows a subsequent memory effect (SME) in the medial temporal lobe (MTL). **(a)** Raster plot for 5 example participants with EEG from hippocampal electrode pairs aligned to time of word presentation. Same participants as the first 5 shown in Sakon & Kahana 2021, Fig. 4b. Each dot represents the start time of a single detected ripples. Vertical gray lines denote the 1.6 s onscreen period for each word, and purple horizontal lines divide participants. We define a trial as a recording from a single bipolar pair during the presentation of a single word. **(b)** Ripple peri-stimulus time histograms (PSTH) averaged across all participants with bipolar electrode pairs localized to hippocampal subfields CA1 or CA3/DG, AMY, or ENTPHC. Each plot displays trials broken into words subsequently recalled or not recalled during the retrieval period. Averages and standard error (SE) bands are from a separate mixed model calculated at each time bin (Eq. 1). When combining across all (recalled and not recalled) words in these four regions, CA1 and CA3/DG show a significant rise in ripples after word presentation compared to baseline (CA1, *β* = 0.019±0.0067, *P* = 0.017; CA3/DG, *β* = 0.017±0.0075, *P* = 0.047), while AMY and ENT/PHC do not (AMY, *β* = 0.0030±0.0066, *P* = 0.78; ENT/PHC, *β* = −0.017±0.0084, *P* = 0.19; each FDR-corrected across 4 tests of Eq. 2)). Significance of mixed model assessing ripple rates between words subsequently recalled vs. not recalled (Eq. 3): CA1, *β* = −0.0029±0.0071, *P* = 0.69; CA3/DG, *β* = −0.0056±0.010, *P* = 0.26; AMY, *β* = 0.0030±0.0066, *P* = 0.69; ENTPHC, *β* = −0.012±0.0064, *P* = 0.69; each FDR-corrected across 4 tests of Eq. 3). Same test for held out participants only: CA1, *β* = 0.0044±0.0095, *P* = 0.65; CA3/DG, *β* = 0.013±0.016, *P* = 0.57; AMY, *β* = 0.0086±0.0074, *P* = 0.57; ENTPHC, *β* = −0.0058±0.0073, *P* = 0.57. **(c)** PSTH for high frequency activity (HFA) using the frequency range 64-178 Hz. HFA is z-scored for each session by averaging across trials and time bins and normalizing with the standard deviation across trials. Error bands are SE from a separate mixed model calculated at each time bin (Eq. 1). Significance of mixed model assessing HFA between words subsequently recalled vs. not recalled (Eq. 3): CA1, *β* = 0.10±0.022, *P* = 1.5 × 10^-5^; CA3/DG, *β* = 0.11±0.028, *P* = 8.5 × 10^-5^; AMY, *β* = 0.16±0.032, *P* = 2.1 × 10^-6^; ENT/PHC, *β* = 0.10±0.030, *P* = 6.1 × 10^-4^ (each FDR-corrected across 4 tests of Eq. 3). Same test for held out participants only: CA1, *β* = 0.11±0.027, *P* = 5.9 × 10^-5^; CA3/DG, *β* = 0.14±0.036, *P* = 1.5 × 10^-4^; AMY, *β* = 0.20±0.035, *P* = 7.3 × 10^-8^; ENTPHC, *β* = 0.11±0.037, *P* = 3.7 × 10^-3^ (each FDR-corrected across 4 tests of Eq. 3).

### High frequency activity (HFA)

We calculate HFA by averaging oscillatory power extracted using Morlet wavelets at 10 logarithmically-spaced frequencies from 64-178 Hz, with the lower bound as in previous HFA work (*7, 16*) and the upper bound the same as for the ripple detector. To measure powers, we use the following procedure using the bipolar-referenced iEEG from each trial from 1 s before word presentation until 2.6 s after word presentation. This window includes a 0.3 s buffer on both sides to avoid edge effects during Morlet transform and 0.7 s (comprised of the inter-trial interval) both before and after word presentation to incorporate as part of the normalization procedure. The signal is then Butterworth filtered from 118-122 Hz and high-pass filtered from 0.5 Hz. A Morlet wavelet transform (using PTSA, see notebooks 5 and 6 on https://github.com/pennmem/CMLWorkshop) is done for each of the 10 frequencies (64.0, 71.7, 80.3, 90.0, 100.8, 113.0, 126.6, 141.8, 158.9, and 178), the buffers are removed, and the log of each value is taken. Next, we resample to 100 ms bins, which leaves us with a FREQUENCY X WORD X CHANNEL X 30 BIN array. We then z-score this array by subtracting the average across words and bins, and dividing by the standard deviation across words after averaging across bins. Finally, we average across the 10 frequencies to arrive at a final HFA value for each WORD X CHANNEL X BIN.

To make the fairest comparison between HFA and ripples, we use the exact same set of trials as selected by our criteria for the ripple detection algorithm. That is, the same word presentations recorded in the same channels (note the identical trial counts in **Fig 2b-c**).

### Anatomical localization

Localization of contacts is identical to previous work (*4*). Briefly, pre-implant structural T1- and T2-weighted MRI scans were used to define the anatomical regions for each participant in addition to a post-implant CT scan to localize electrodes in the participant brain, which were coregistered using Advanced Normalization Tools (*25*). The point source of iEEG for bipolar electrode pairs is considered to be the midpoint between adjacent electrode contacts. Center to center electrode spacing was between 3-10 mm as chosen by the neurosurgical teams for medical reasons.

Similar to our previous work (*4*), we split channels localized to hippocampus into two groups, CA1 and CA3/DG, since we have sufficient sample size to test our hypotheses in each. However, since we use the midpoint of bipolar electrode pairs for signal localization (hippocampal pairs are 3-6 mm apart as only stereo-EEG depth electrodes reach hippocampus), and considering an estimated 350,000 neurons contribute to macroelectrode LFP (*4*), many of the channels are likely to reflect ripples crossing subfields.

Bipolar electrode pairs in hippocampal subfields CA1 and dentate gyrus (DG) were localized using a combination of neuroradiologist labels (Joel M. Stein and Sandhitsu Das, Penn Medicine) and the automated segmentation of hippocampal subfields (ASHS) technique utilizing the T2 scan (*26*). However, we label the DG pairs as CA3/DG due to the difficulty in delineating these regions. Sites localized to CA3 are not included in this group as ASHS achieves poor classification of this subfield compared to CA1 and DG (*26*)), and because of its relatively small volume (*∼*15x fewer channels are localized to CA3 than DG).

We also analyze electrode pairs in non-hippocampal cortical regions, which include entorhinal (ENT), parahippocampal (PHC), and perirhinal cortex and amygdala. We used a combination of neuroradiologist labels and an automated segmentation pipeline combining whole-brain cortical reconstructions from the T1 scan in Freesurfer (*27*), an energy minimization algorithm to snap electrodes to the cortical surface (*28*), and boundaries and labels from the Desikan-Killiany-Tourville cortical parcellation protocol (*29, 30*)).

### Plots and binning

Raster plots are formed by aligning the iEEG to the time of word presentation and plotting the time of the beginning of each detected ripple. Peri-stimulus time histograms (PSTHs) are formed by binning ripples (100 ms bins) and averaging the raster plots across participants after separating words into groups (e.g. subsequently recalled vs. not recalled words). For visualization only, these PSTHs are triangle smoothed using a 5-bin window (*4, 5*) and a separate linear mixed model with sessions nested in participants is run at each bin to calculate the mean and error bars (SE) (**Eq. 1**). Ripple rates are the frequency in Hz. within each bin.

The default analysis window used to assess the ripple subsequent memory effect (SME) and the subsequent clustering effect (SCE) throughout the paper is 0.1 to 1.7 s from beginning of word presentation. We offset 0.1 s from time on screen to account for latencies from the time of presentation until signals reach MTL circuits (*31*). The analysis window for HFA is from 0.4 to 1.1 s after word presentation. These windows are based on pilot analyses done on the first half of the data and pre-registered on the Open Science Framework (OSF, https://osf.io/e98qp). We also report statistics for the SME from 0.4 to 1.1 s as a comparison to the window used for HFA. To measure pre-retrieval effect (PRE) ripples during the retrieval period we use the window from −1.1 to −0.1 s prior to recall vocalization (*4*). To assess the rise in ripples after word onset we use −0.7 to 0.1 s as for the baseline ripple rate and 0.1 to 0.9 s as an equally-sized window locked to stimulus onset.

### Clustering

When participants correctly recall a series of words during the retrieval period, the order of word recall provides a window into the organization of their memory. For categorized free recall, as participants transition from one recall to the next, we expect them to cluster recalls based on semantic and/or temporal relationships between words on the list. As explained in the Free Recall task section above, each 12-word list in this task had words drawn from 3 categories, with the 4 words from the same category presented in non-contiguous pairs. This setup provides three distinct forms of clustering between consecutive recalls: adjacent semantic (20% of recalls lead to this transition), remote semantic (20%), and adjacent non-semantic (3%) (examples given in **Fig. 1b-c**). Adjacent semantic are two words from the same category shown as a consecutive pair during encoding while remote semantic are two words from the same category from pairs separated by other words. Adjacent, non-semantic transitions were not analyzed due to their small sample size. Recalls that do not lead to clustering include remote unclustered (17%), where consecutive words were neither from the same category or shown back-to-back, and dead ends (26%), which are the last recall that do not lead to a subsequent recall. The remaining recalls were those that led to intrusions or repeats (14%).

For the SCE contrast we pool clustered and unclustered recalls in **Fig. 3b**, and for the SME contrast we pool unclustered recalls in **Fig. 3c**. However, in the caption for the SCE contrast, we also provide statistics for pairwise models between each of the clustering types (adjacent semantic and remote semantic) vs. unclustered recalls. And in the caption for the SME contrast, we also provide statistics for pairwise models between each of the unclustered types (remote unclustered and dead ends) vs. not recalled words.

**Figure 3.**
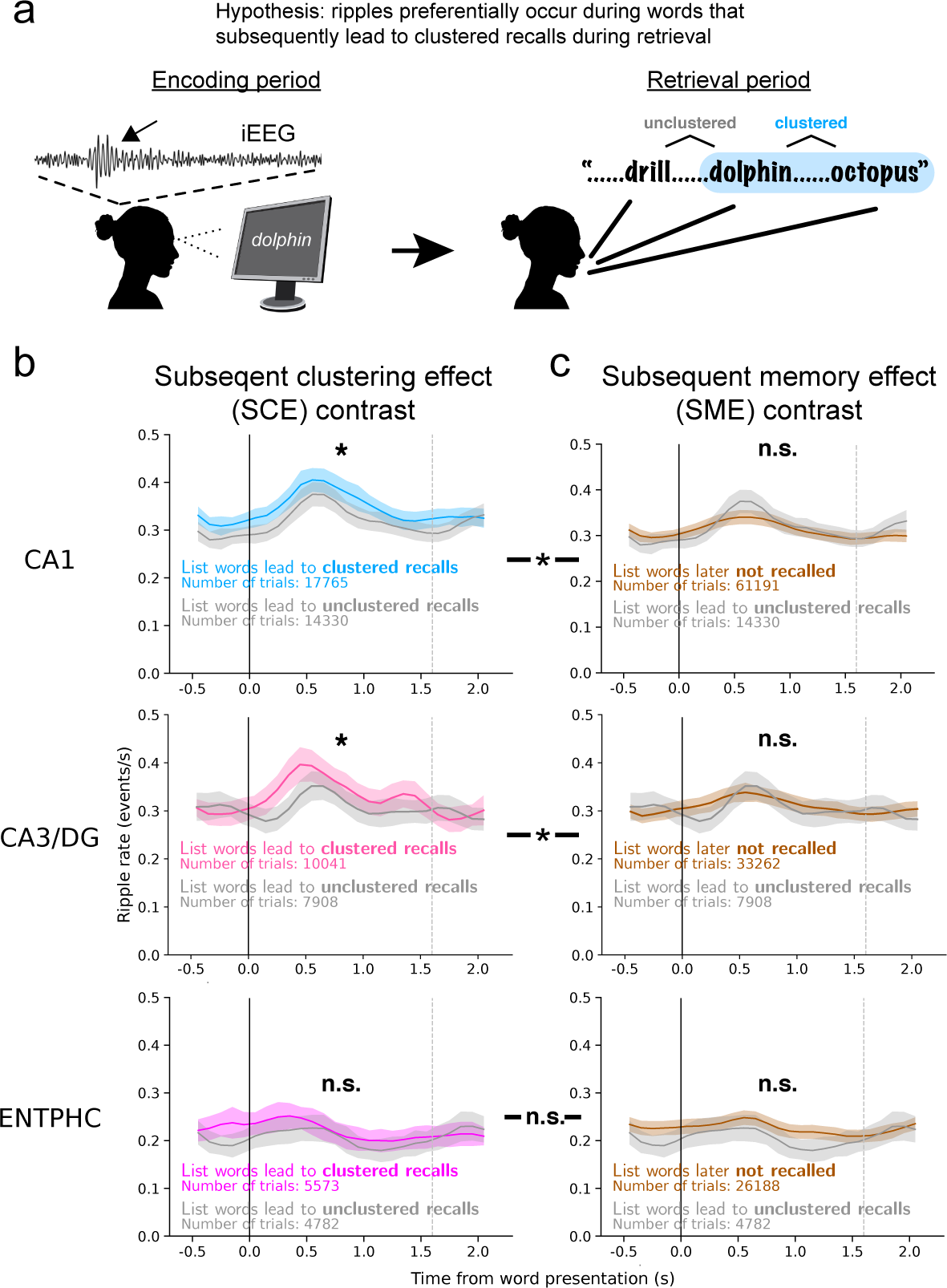
Hippocampal ripples signal a subsequent clustering effect (SCE). **(a)** Diagram of the SCE. When words are recalled during the retrieval period (right) we examine the relationships between the recall order to identify semantic or temporal relationships. Using the example list shown throughout the manuscript (see Fig. 1b), dolphin and octopus are adjacent semantic as they were a pair shown back-to-back and are from the same semantic category. We then measure ripples during the encoding period (left) when dolphin was presented as this was the word that led to the subsequent transition (or clustering) between recalls during retrieval. **(b)** Ripples rates grouped by clustering category for CA1, CA3/DG, and ENTPHC sites. Each plot shows words that lead to subsequent clustering (adjacent semantic and remote semantic) vs. those that do not (remote unclustered combined with dead ends). Significance of mixed model comparing clustered vs. unclustered groups for each region: CA1, *β* = 0.022±0.0080, *P* = 0.015; CA3/DG, *β* = 0.031±0.012, *P* = 0.017; ENTPHC, *β* = 0.015±0.0088, *P* = 0.089 (each FDR-corrected across 3 tests of Eq. 3). Same test for held out participants only: CA1, *β* = 0.012±0.0091, *P* = 0.30; CA3/DG, *β* = 0.024±0.019, *P* = 0.30; ENTPHC, *β* = 0.012±0.011, *P* = 0.31 (each FDR-corrected across 3 tests of Eq. 3). **(c)** Each plot shows a breakdown of words subsequently recalled but without clustering (remote unclustered and dead ends) vs. those not recalled. Significance of mixed model comparing these groups for each region: CA1, *β* = −0.013±0.0079, *P* = 0.098; CA3/DG, *β* = −0.019±0.018, *P* = 0.079; ENTPHC, *β* = −0.020±0.0081, *P* = 0.040 (each FDR-corrected across 3 tests of Eq. 3). Same test for held out participants only: CA1, *β* = −0.0036±0.010, *P* = 0.73; CA3/DG, *β* = −0.021±0.014, *P* = 0.40; ENTPHC, *β* = −0.011±0.010, *P* = 0.40 (each FDR-corrected across 3 tests of Eq. 3). For all plots vertical black and gray lines denote word presentation onset and offset and error bands are SE from a separate mixed model calculated at each time bin (Eq. 1). Asterisks between the left and right plots indicate the SCE is significantly greater than the SME for that region (Eq. 4, see main text).

### Held out data and pre-registration

The large size of our dataset allowed us to set aside *∼*35% of trials in order to come up with initial figures and hypotheses that can then be confirmed with the entire dataset. That is, after creating a raster plot to ensure all data is in usable form after the data-cleaning steps outlined in Ripple Detection above, we used a random kernel to select a subset of participants comprising 35% of hippocampal trials. Once we set our initial analysis parameters and figures based on this exploratory 35% of data, we registered them along with hypotheses based on these figures on the Open Science Framework (https://osf.io/e98qp), which also contains specific details on the randomization and sampling plan. Here we present the statistics and figures for the entire dataset based on the analysis parameters defined in this pre-registration.

### Equations

Linear mixed effects models are run using the function MixedLM in the python package statsmodels with restricted maximum likelihood and Nelder-Mead optimization with a maximum of 2000 iterations. The following equations are written in pseudocode of the inputs to statsmodels. Statistics are presented as: *β ± SE, P − value*, where *β* is the coefficient being tested and *SE* is the standard error of the coefficient being fit. For all comparisons the first group takes the indicator value 1 and the second takes 0 in the model. For example, clustered vs. unclustered trials are assigned 1 and 0, meaning if clustered is greater the coefficient will be positive.

We use mixed effects models to plot the mean and standard error of ripple rates for all peri-stimulus time histograms (PSTHs). For a given group of trials, a separate mixed effect model is run at each 100 ms bin:

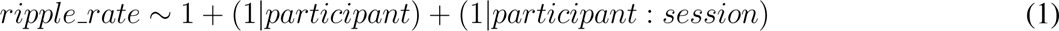

where (1*|participant*) is a random intercept and slope for each participant, (1*|participant* : *session*) is a random intercept and slope for each session nested in each participant, and *ripple rate* is the average ripple rate in that bin for a given trial. The solved coefficient and its standard error (SE) are used to plot the mean *±* SE at each bin (after a 5-point triangle smooth of the means). Plotting the average ripple rates across trials looks similar, but plots using the mixed effects mean have the advantage of 1) giving a better estimate of the population mean after accounting for inter-subject sample sizes and differences in ripple rates and 2) providing a more accurate visualization of the statistical fits used to compare groups of trials in the following equations.

We assess the rise in ripple rates after word presentation using the following model:

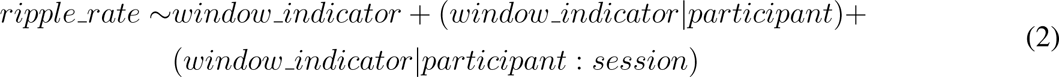

where *window indicator* is an indicator variable with value 1 for the window from 0.1 to 0.9 s aligned to word presentation and value 0 for the window from −0.7 to 0.1 s aligned to word presentation, (*window indicator|participant*) are random intercepts and slopes for each participant, (*window indicator|participant* : *session*) are random intercepts and slopes for sessions nested in each participant, and *ripple rate* is the average ripple rate within the given window for that trial. We use −0.7 to 0.1 s for the baseline ripple rate since the minimum inter-stimulus interval is 0.75 s and MTL ripples have not been shown to occur until *>*0.1 s after stimulus presentation (*5, 7, 8*). To capture the rise in ripples we use an adjacent period of equal duration from 0.1 to 0.9 s that encompasses the peak ripples rates clearly seen in the raster (**Fig. 2a**) and PVTHs (**Fig. 2b-c**). The null hypothesis is no difference between ripple rates for the two windows.

To test the hypothesis that ripples rates increase during words that are subsequently recalled vs. subsequently not recalled (**Fig. 2b & 3b**), we use the linear mixed effects model:

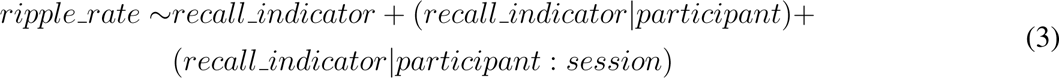

where *recall indicator* is an indicator variable with value 1 for words subsequently recalled and 0 for those that are not and *ripple rate* is the average ripple rate for each trial from 0.1 to 1.7 s following word presentation. Random intercepts and slopes for sessions nested in participants follow the same structure as **Eq. 2**. The null hypothesis is no difference between ripple rates on words that are subsequently remembered v. subsequently not recalled.

We use the same model for comparisons between groups, such as words that subsequently lead to clustered recalls vs. unclustered recalls **Fig. 3**). In this case, instead of recall indicator, the predictor indicates if a recalled word subsequently leads to clustering or not (e.g. subsequently clustered vs. unclustered recalls, **Fig. 3b**). We also use this model to compare SMEs for HFA from 0.4-1.1 s following word presentation and as a comparison SMEs for ripples from 0.4-1.1 s.

We compare the SCE and the SME directly in the same model:

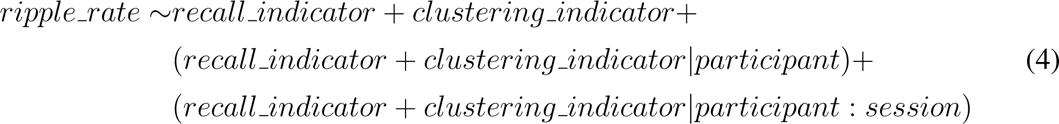

where *recall indicator* and *clustering indicator* are the same as defined below **Eq. 3**. Random intercepts and slopes for sessions nested in participants follow the same structure as **Eq. 2**. Note that the equation adds the two indicator variables instead of multiplying them because it is not possible to have a trial coded with 0 for recall and 1 for clustering. The null hypothesis is no difference in ripples between clustered and unclustered words after taking into account recalled vs. not recalled words.

We hypothesize that participants that recall more words will show a bigger ripple subsequent clustering effect (SCE), in which words that subsequently are recalled and lead to clustering will have more ripples than words that are subsequently recalled and do not lead to clustering. To test this relationship we use the linear mixed effects model:

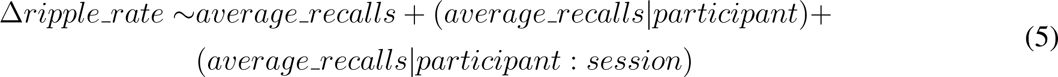

where *average recalls* is the average number of recalls per 12-word list for the participant and Δ*ripple rate* is the average difference in ripple rate from 0.1 to 1.7 s following word presentation for subsequently clustered (i.e. adjacent semantic and remote semantic trials) vs. unclustered (i.e. remote unclustered and dead ends) words. Random intercepts and slopes for sessions nested in participants follow the same structure as **Eq. 2**. The null hypothesis is that SCE does not relate to memory performance. For **Fig. 4**, including this model and the following one, we only include patients with at least 20 clustered and 20 unclustered trials for the full dataset analyses and at least 10 of each for the held out data analyses.

**Figure 4.**
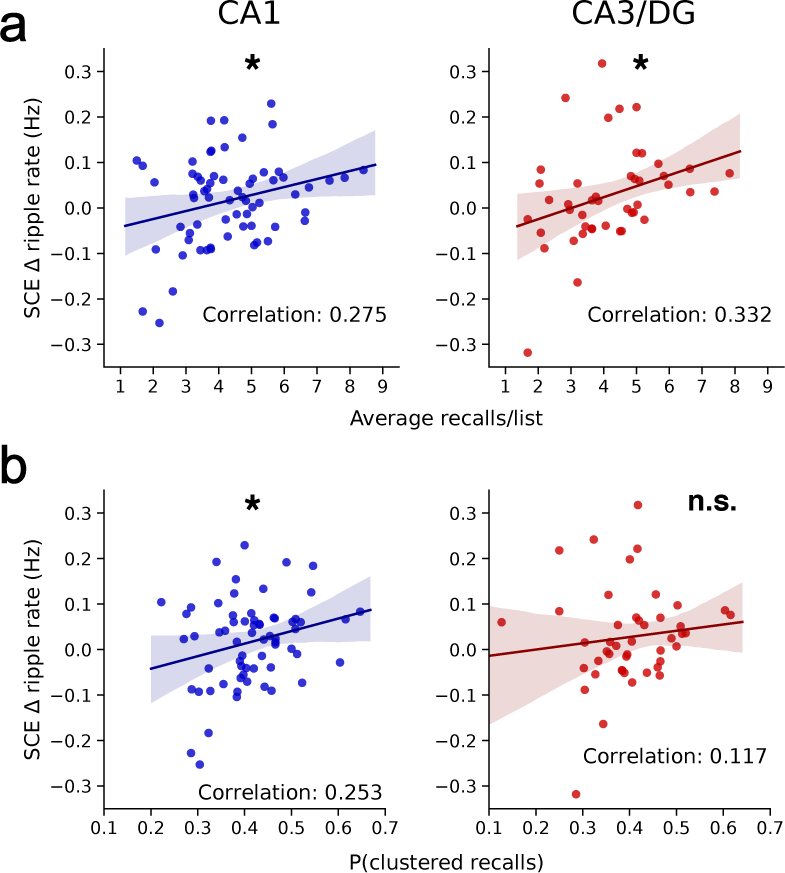
The ripple subsequent clustering effect (SCE) relates to memory performance. **(a)** For each participant, we relate the change in ripple rate during word presentation between recalled words that lead to subsequent clustering (adjacent semantic and remote semantic) vs. no clustering (remote unclustered and dead ends) to the average number of words recalled on each list by that participant. Significance of mixed model comparing this change in ripple rate SCE vs. the average recalls per list: CA1, *β* = 0.015±0.0056, *P* = 0.0089; CA3/DG, *β* = 0.023±0.0.0084, *P* = 0.0089 (each FDR-corrected across 2 tests of Eq. 5). Same test for held out participants only: CA1, *β* = 0.016±0.0071, *P* = 0.028; CA3/DG, *β* = 0.036±0.011, *P* = 0.0015 (each FDR-corrected across 2 tests of Eq. 3). **(b)** For each participant, we relate the same change in ripple rate from **a** to the average proportion of clustered recalls (i.e. recalls that are adjacent semantic and remote semantic out of all recalls). Significance of mixed model comparing the SCE vs. the proportion of clustered recalls: CA1, *β* = 0.25±0.11, *P* = 0.046; CA3/DG, *β* = 0.26±0.18, *P* = 0.14 (each FDR-corrected across 2 tests of Eq. 6). Same test for held out participants only: CA1, *β* = 0.39±0.15, *P* = 0.0096; CA3/DG, *β* = 0.54±0.19, *P* = 0.0096 (each FDR-corrected across 2 tests of Eq. 3). Both plots in this figure and the whole dataset models use only patients with at least 20 clustered and 20 unclustered trials; held out models require at least 10 of each.

We also compare the SCE Δ*ripple rate* with the amount of clustering at the participant-level using a similar linear mixed-effects model:

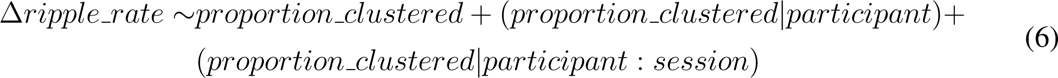

where *proportion clustered* is the combined number of words that lead to adjacent semantic and remote semantic trials divided by the total number of words recalled for each participant. Random intercepts and slopes for sessions nested in participants follow the same structure as **Eq. 2**. The null hypothesis is that SCE does not relate to the amount participants recall words via clustering.

Next we investigate the hypothesis that a ripple during the first pair of words from a category (*X*_1_*_−_*_2_) will make it more likely to see reinstatement–and therefore a ripple (*4*)–during the second pair of words from a category (*X*_3_*_−_*_4_). As a result, we expect likelier recall of *X*_3_*_−_*_4_ if a ripple occurs during *X*_1_*_−_*_2_, and even likelier recall if a ripple occurs during both pairs. To test this hypothesis we use the linear mixed-effect model:

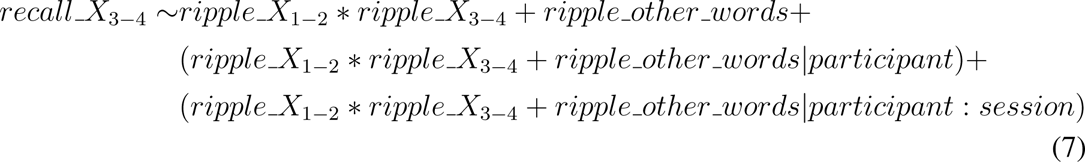

where *recall X*_3_*_−_*_4_ indicates if a participant recalled a word from *X*_3_*_−_*_4_, *ripple X*_1_*_−_*_2_ indicates a ripple occurred during *X*_1_*_−_*_2_, *ripple X*_3_*_−_*_4_ indicates a ripple occurred during *X*_3_*_−_*_4_, and *ripple other words* is the ripple rate for the remaining (eight) words on the list not from that category. The * indicates separate coefficients are calculated for each term and the interaction. Random intercepts and slopes for sessions nested in participants follow the same structure as **Eq. 2**. The null hypotheses are that 1) recall of a word from *X*_3_*_−_*_4_ is not more likely if a ripple occurs during *X*_1_*_−_*_2_ (the coefficient for *ripple X*_1_*_−_*_2_) and 2) recall of a word from *X*_3_*_−_*_4_ is not more likely if a ripple occurs during both *X*_1_*_−_*_2_ and *X*_3_*_−_*_4_ (the coefficient for the interaction *ripple X*_1_*_−_*_2_ : *ripple X*_3_*_−_*_4_).

As a control, we use the same model as above to predict *X*_1_*_−_*_2_ recalls (instead of *X*_3_*_−_*_4_ recalls). The null hypothesis is recall of words from *X*_1_*_−_*_2_ is not more likely if a ripple occurs during both *X*_1_*_−_*_2_ and *X*_3_*_−_*_4_.

Finally, we test the hypothesis that a ripple during encoding of a word combined with a ripple in the PRE window during its subsequent recall will increase the likelihood that word leads to clustering. To test this hypothesis we use the linear mixed effects model:

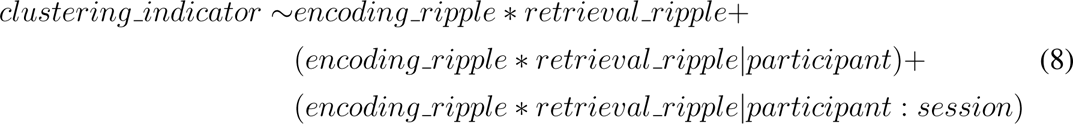

where *clustering indicator* is 1 if a recalled word leads to clustering and 0 if not (i.e. remote unclustered or dead end), *encoding ripple* is an indicator variable with the value 1 if *≥*1 ripple occurred in the window from 0.1 to 1.7 s after word presentation, and *retrieval ripple* is an indicator variable with the value 1 if *≥*1 ripple occurred in the window from −1.1 to −0.1 s aligned to vocalization of the word during retrieval. The * indicates separate coefficients are calculated for each term and the interaction. Random intercepts and slopes for sessions nested in participants follow the same structure as **Eq. 2**. The null hypothesis is no increase in clustering when a ripple occurs during encoding of a word and prior to its subsequent recall.

## Results

### Hippocampal ripples do not exhibit a subsequent memory effect (SME)

To clarify the relation between ripples and memory encoding we align hippocampal recordings (**Fig. 1e**) to the onset of word presentation during the study phase of a categorized, delayed free recall task (**Fig. 1b**), in which participants view a list of words and subsequently recall as many as possible after a distractor period. We use an algorithm recently shown to isolate ripples in human hippocampus and surrounding MTL during memory retrieval (*4, 5*) (*Methods*). A raster of ripples from five sample participants illustrates an encoding-related rise in ripples occurring *∼*0.5 seconds after word onset (each row in **Fig. 2a** represents a word presentation recorded on a single channel, and each dot represents the start time of a single ripple). Measuring the rate of ripples across all trials after word presentation as compared with the baseline rate prior to word presentation, both hippocampal subfields CA1 (163 sessions from 86 participants) and CA3/dentate gyrus (CA3/DG: 117 from 59; 54 overlapping participants with CA1) show a significant rise (**Fig. 2b**). However, amygdala (AMY; 104 sessions from 50 participants; 33 overlapping with either hippocampal subfield) and entorhinal/parahippocampal cortex (ENT-PHC, 96 from 52; 33 overlapping with either hippocampal subfield and 28 overlapping with AMY) fail to show a significant increase in ripples after word presentation (**Fig. 2b**). These findings accord with prior work where hippocampal ripples increase hundreds of ms after presentation of face or place stimuli (*5, 7, 8*).

For our first test of encoding we ask if ripples show an SME. Once again separately investigating both hippocampal subfields CA1 and CA3/DG, we average across participants to create peri-stimulus time histograms (PSTHs) for both subsequently recalled and not recalled words. We find only a modest difference in ripple rates between these groups beginning *∼*0.5 seconds after word presentation in both regions (**Fig. 2b**). Although the dataset is adequately powered to find a ripple SME in each region (power*>*0.97 using effect sizes for SMEs reported using HFA (*21*) and ripple (*7*) detectors, Methods), we find no significant difference between ripple rates during word presentation of subsequently recalled vs. not recalled words for either CA1 or CA3/DG (**Fig. 2b**; **Eq. 3**). Meanwhile, both AMY and ENTPHC show overall lower ripple rates than the hippocampal subfields and also fail to show a ripple SME (**Fig. 2b**).

Considering that previous studies find strong HFA SMEs in the hippocampus and neighboring MTL subregions (*16, 32*) we apply a high frequency activity (HFA) detector on the same trials as in the ripple analysis reported above. Measuring HFA in a frequency range almost completely overlapping that of our ripple detector, we find a clear HFA SME in all MTL subregions (**Fig. 2c**). Using the same linear mixed effects model as with ripples, the HFA SME is significant for CA1, CA3/DG, AMY, and ENTPHC (**Fig. 2c**; **Eq. 1**). Notably, when assessing the ripple SME with this model using a smaller time window that matches the significant range for the HFA SME (0.4-1.1 s), all four regions still fail to show a significant ripple SME (*P>*0.30, each FDR-corrected across 4 tests of Eq. 3). In sum, HFA exhibits an SME across the MTL, while the ripple detector does not for any MTL region. The overlapping frequency range between the detectors suggests that this difference comes from the extra processing steps in the ripple detection algorithm (Discussion).

### Hippocampal ripples exhibit a subsequent clustering effect (SCE)

The SME contrast fails to take advantage of the rich behavioral structure of the categorized free recall task (**Fig. 1b-c**). Specifically, the order in which people free recall recently studied items reveals information about the organization of memory. When participants strongly bind items to their encoding context, which includes both temporal and semantic information (*33*), they tend to retrieve clusters of temporally and semantically similar items (*22, 34*). Our previous work showed an increase in hippocampal ripples just prior to participants recalling a cluster of related words, suggesting that ripples signal the reinstatement of context (*4*). Here, we hypothesize that ripples might also signal contextual reinstatement during encoding. If this is true, an increase in ripples during initial presentation of a word predicts that word will subsequently lead to clustering during retrieval (**Fig. 3a**). We refer to this phenomenon as a subsequent clustering effect (SCE) (*21*).

In categorized free recall, transitions between clustered recalls neatly divide into a handful of groups (**Fig 1c**). Referring to the example words in **Fig 1c**: **adjacent semantic** indicates two words from the same categorical pair recalled consecutively, e.g. dolphin and octopus (22% of recalls); **remote semantic** indicates two words from the same category but not the same pair recalled consecutively, e.g. dolphin and fish (24% of recalls); **remote unclustered** indicates two words from different categories that are not presented back-to-back recalled consecutively, e.g. dolphin and pliers (22% of recalls); and **dead end** indicates the last recall from each list, which therefore does not transition to another recall (15% of recalls). Adjacent unclustered, in which participants recall words that appear back-to-back from different categories, are rare (4%) so we do not analyze this type further, while the remaining recalls are incorrect (12%) or repeats (2%). We then measure ripples during the presentation of the first word in each transition pair (except for dead ends, where there is no transition) and compare ripple rates between transition types.

Measuring the average ripple rates between these behaviorally-defined groups reveals clear evidence that hippocampal ripples exhibit an SCE. In particular, testing the ripple rates of words that lead to subsequent clustering (adjacent semantic and remote semantic) vs. those that are subsequently recalled but do not lead to clustering (remote unclustered and dead ends) yields a significant difference in both CA1 and CA3/DG, but not in ENTPHC (**Fig 3b**, **Eq. 3**). When making comparisons between the individual categories in the clustering group (i.e. adjacent semantic and remote semantic), each of these also show significantly more ripples during their presentation compared to unclustered recalls for both CA1 and CA3/DG (p*≤*0.031, FDR-corrected across six individual tests, **Eq. 3**) but not ENTPHC (p*≥*0.18, FDR-corrected across six individual tests, **Eq. 3**).

The previous contrasts isolate clustering as we compare words that subsequently lead to clustering vs. those words that are still recalled but do not lead to the subsequent semantic or temporal transitions that hallmark context reinstatement. In a similar manner, we can isolate the ripple SME by contrasting recalled words that do not lead to clustering vs. words not recalled. Using this contrast, we find no evidence of an SME in CA1, CA3/DG, or ENTPHC (**Fig. 3c**, **Eq. 3**). In fact, all three regions have negative coefficients for Eq. 3, indicating a ‘reverse’ ripple SME as fewer ripples occur during words subsequently leading to unclustered recalls than words not recalled, although only ENTPHC shows a significant difference (**Fig. 3c**, **Eq. 3**). Further, when making comparisons between the individual categories in the recalled but not clustered group (i.e. remote semantic and dead ends) vs. words not recalled, each of these also show no significant difference in ripples rates for CA1, CA3/DG, or ENTPHC (*P>*0.11, FDR-corrected across six individual tests, **Eq. 3**).

To directly compare the ripple SCE and SME, we contrast subsequently clustered words with unclustered words in the same statistical test (**Eq. 4**). This model allows us to measure the significance of the SCE after taking into account ripples during the remaining subsequently recalled words. Both CA1 and CA3/DG (*P<*0.025), but not ENTPHC (*P* = 0.14), have a significant positive coefficient for the SCE factor (FDR-corrected across three tests of **Eq. 4**), indicating an increase in ripples specific to subsequently clustered words. All three regions show a negative coefficient for the SME, with CA3/DG and ENTPHC significant (*P* = .038 for each; *P* = 0.069 for CA1, FDR-corrected across three tests of **Eq. 4**), indicating fewer ripples during subsequent unclustered recalls compared to words not recalled. Therefore, hippocampal ripples rise specifically during words that subsequently lead to clustered recalls.

Considering that HFA shows an SME while ripples do not (**Fig. 2**), does HFA also reflect an SCE? Our expectation is that due to the highly overlapping frequency ranges of these two detectors, HFA will pick up both the SCE from ripples in addition to the SME we have already shown. We validate this prediction. Comparing HFA during subsequently recalled words that lead to clustering vs. subsequently recalled words that do not, CA1 and CA3/DG show significantly stronger HFA (*P<*0.020), while ENTPHC does not (p=0.33, each FDR-corrected across three tests of **Eq. 3**). Meanwhile, all three brain regions show an HFA SME when comparing not recalled words to words subsequently recalled but not clustered (*P<*0.023, FDR-corrected across three tests of **Eq. 3**). Finally, when comparing HFA SCE and SME in the same model, CA1 and CA3/DG each show significant rises in HFA for both the SCE (*P<*0.033) and the SME (*P<*0.041, FDR-corrected across three tests of **Eq. 4**). ENTPHC only shows a significant increase in HFA for the SME (*P* = 0.036, FDR-corrected) but not for the SCE (*P* = 0.28). In conclusion, hippocampal HFA can reflect either subsequent clustering or subsequent memory during word encoding, unlike hippocampal ripples which specifically reflect encoding of words that lead to subsequent clustering of recalls.

### The hippocampal ripple SCE is associated with better memory and increased clustering

Next, we ask if the hippocampal SCE shown in **Fig. 3** correlates with participant behavior. Measuring the SCE for each individual participant as the difference in ripples during words that lead to subsequent clustering vs. recalled words that do not lead to clustering, we compare this change to the average number of recalls for that person per list. Participants that recall more words display a significantly larger ripple SCE in both CA1 and CA3/DG (**Fig. 4a**, **Eq. 5**). Therefore, the hippocampal ripple SCE predicts superior memory across participants.

Does the hippocampal ripple SCE also predict clustering of recalls? Contrasting the ripple SCE with the proportion of recalls that lead to subsequent clustering out of all recalls, both CA1 and CA3/DG show a positive correlation although only CA1 is significant (**Fig. 4b**, **Eq. 6**). Therefore, participants with a larger hippocampal ripple SCE both remember more words and more frequently recall them via clustering.

### Hippocampal ripples during both the first and second pair of words from a category lead to improved recall of the second pair

On each list in the free recall task two pairs of words from the same semantic category appear: one in the first half of the list and another in the second half (with the constraint that pairs from the same category are never shown back-to-back) (**Fig. 1b**). This task structure allows us to investigate if ripples occurring as participants encode the first pair of words from a given category (*X*_1_*_−_*_2_) influence memory for the second pair (*X*_3_*_−_*_4_) despite the intervening word presentations. Considering the SCE results (**Fig. 3**), in which increased ripples during word presentation predict the word will subsequently lead to context reinstatement (and therefore clustering) during the retrieval period, we hypothesize that ripples during *X*_1_*_−_*_2_ might also lead to context reinstatement during the presentation of words *X*_3_*_−_*_4_. And if such context reinstatement manifests during the presentation of words *X*_3_*_−_*_4_, we anticipate likelier subsequent recall of these words (**Fig. 5a**).

**Figure 5.**
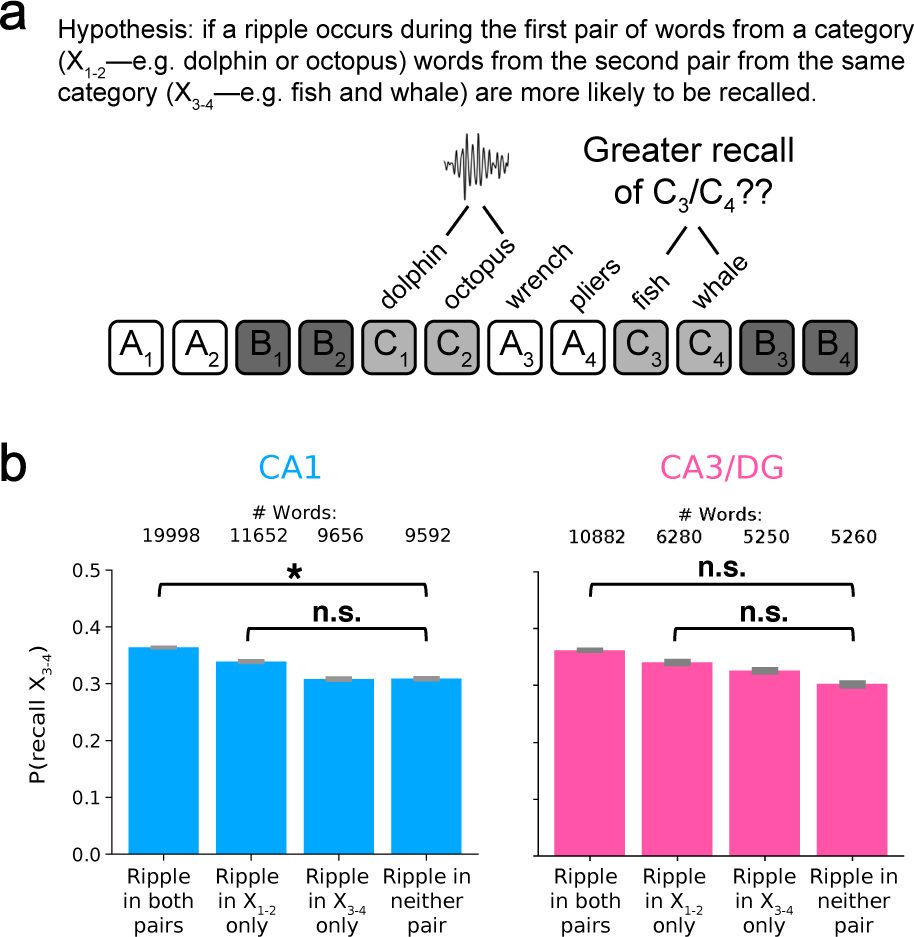
Presence of hippocampal ripples during initial category presentation leads to better recall of words from the same category. **(a)** Diagram of hypothesis that ripples during presentation of words from a category will increase likelihood of recalling subsequently presented words from same category. **(b)** Accuracy of recall for the second pair of words from a category (*X*_3_*_−_*_4_) when a ripple occurs during either of the first pair of words from a category (*X*_1_*_−_*_2_), either of the second pair of words from a category (*X*_1_*_−_*_2_), both, or neither. The number of total words for each of these pools is indicated above the bars. Error bars are SE of proportions. Significance of mixed model term assessing the impact on accuracy for *X*_3_*_−_*_4_ based on the presence of ripples during *X*_1_*_−_*_2_: CA1, *β* = −0.012±0.0063, *P* = 0.12; CA3/DG, *β* = −0.0059±0.0087, *P* = 0.50 (each FDR-corrected across two tests of Eq. 7). Significance of mixed model term assessing the impact on accuracy for *X*_3_*_−_*_4_ based on the presence of ripples during both *X*_1_*_−_*_2_ and *X*_3_*_−_*_4_: CA1, *β* = 0.023±0.0083, *P* = 0.010; CA3/DG, *β* = 0.012±0.011, *P* = 0.30 (each FDR-corrected across two tests of Eq. 7).

To test this hypothesis, we measure the accuracy of *X*_3_*_−_*_4_ words from each category on each list after assigning the category to one of four pools: 1) those where *≥*1 ripple occurs during the presentation of *X*_1_*_−_*_2_ (but not *X*_3_*_−_*_4_), 2) those where *≥*1 ripple occurs during *X*_3_*_−_*_4_ (but not *X*_1_*_−_*_2_), 3) those where *≥*1 ripple occurs during both *X*_1_*_−_*_2_ and *X*_3_*_−_*_4_, and 4) those where no ripple occurs during either. Averaging within each pool, we find words with a ripple during both *X*_1_*_−_*_2_ and *X*_3_*_−_*_4_ exhibit the highest recall accuracy, followed by lists with ripples only during *X*_1_*_−_*_2_ (**Fig. 5b**).

To evaluate differences in the accuracy of *X*_3_*_−_*_4_ recall among the pools, we create a linear mixed model that takes into account *≥*1 ripple during presentation of *X*_1_*_−_*_2_, *≥*1 ripple during presentation of *X*_3_*_−_*_4_, the interaction of a ripple occurring for both pairs, and also the ripple rate for the remaining (eight) words on the list to remove possible list-level ripple rate effects (**Eq. 7**). This model reveals that CA1 ripples during *X*_1_*_−_*_2_ predict *X*_3_*_−_*_4_ recall, but *only* if a ripple also occurs during *X*_3_*_−_*_4_ (**Fig. 5b**). Thus, if a ripple occurs during both pairs of words from a category, the likelihood of recalling the 2nd pair (*X*_3_*_−_*_4_) increases. However, if a CA1 ripple occurs only during *X*_1_*_−_*_2_, we find no significant difference in recall accuracy of *X*_3_*_−_*_4_. CA3/DG does not show a significant difference for either comparison, even though the effect is in the same direction for better *X*_3_*_−_*_4_ recall when a ripple occurs during both pairs (**Fig. 5b**).

If the increase in *X*_3_*_−_*_4_ recalls comes from *X*_1_*_−_*_2_ ripples leading to context reinstatement and therefore ripples during *X*_3_*_−_*_4_, as opposed to an additive effect where increased ripples during same category words leads to more recalls from that category, we anticipate that ripples during both pairs of words will not improve recall of *X*_1_*_−_*_2_ recalls. Indeed, when ripples occur during both *X*_1_*_−_*_2_ and *X*_3_*_−_*_4_, recall of *X*_1_*_−_*_2_ words does not increase when measuring either CA1 or CA3/DG ripples (p*>*0.37, **Eq. 7**). These findings suggest ripples during early list words promote category reinstatement later in the list, as reflected by ripples occurring for words from the same category later in the list.

### Hippocampal ripples during encoding and retrieval of the same word predict clustering

The previous analysis suggests that ripples can reflect context reinstatement during encoding, where ripples during early list words promote ripples during late list words when the words carry strong semantic relations. Our previous work finds ripples reflect context reinstatement during retrieval, as ripples occur just prior to vocalization of clustered recalls (the pre-retrieval effect (PRE), see Discussion). Here we ask whether clustering emerges specifically when ripples occur during both encoding and retrieval of the same words (**Fig. 6a**).

**Figure 6.**
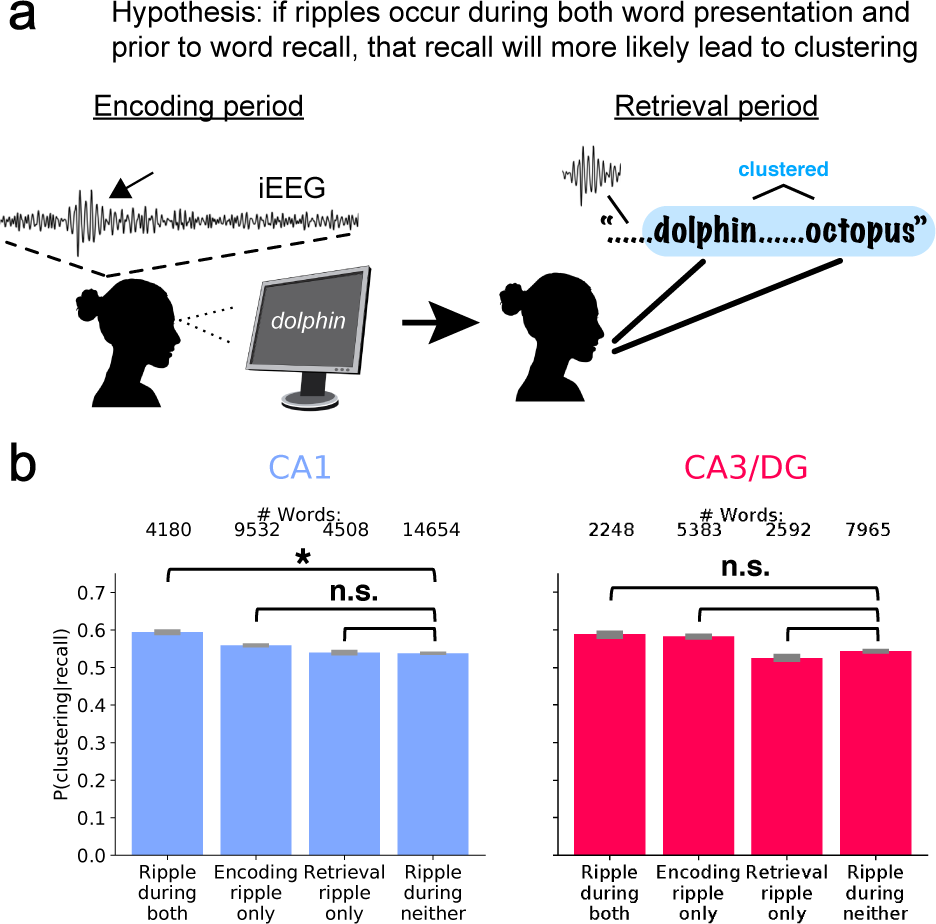
Words with ripples during both word presentation and prior to recall lead to clustering. **(a)** Diagram of hypothesis that clustering arises when ripples occur during both the presentation of and prior to the recall of words. **(b)** Proportion of recalls that lead to clustering conditioned on whether the recalled word has *≥*1 ripple occur during its initial presentation and/or prior to its vocalization. The number of total recalls for each condition is indicated above the bars. Error bars are SE of proportions. Significance of mixed model terms assessing the impact on clustering of the presence of ripples during both word encoding or retrieval: CA1, *β* = 0.037±0.0074, *P* = 9.5 × 10^-7^; CA3/DG, *β* = 0.023±0.017, *P* = 0.17 (each FDR-corrected across 2 tests of **Eq. 8**). P-values for remaining terms are not significant (*P ≥*0.065, each FDR-corrected).

To answer this question, for every recalled word we determine if *≥*1 ripple occurs during its presentation and/or during the PRE window. Assigning each recall to one of four conditions— encoding *±* ripple crossed with retrieval *±* ripple—we assess the proportion of recalls within each condition that lead to clustering. As predicted, recalls with ripples during both encoding and retrieval exhibit the highest clustering rates **Fig. 6b**. Using a linear mixed effects model to assess if ripples during encoding, retrieval, or both lead to clustering, only when CA1 ripples occur in both conditions do we find a significant increase in clustering **Fig. 6b, left**. Ripples measured in CA3/DG, however, do not significantly predict clustering regardless of their presence during encoding, retrieval, or both periods **Fig. 6b, right**.

## Discussion

Measuring medial temporal lobe (MTL) ripples as participants encode and then free recall lists of words, we find that clustering of recalls significantly increases during memory retrieval specifically when hippocampal ripples occur during word presentation. This ripple subsequent clustering effect (SCE) appears more prominently than a ripple subsequent memory effect (SME), specifying a role for ripples in binding items to their semantic and/or temporal associates when forming memories. The magnitude of the hippocampal ripple SCE also aligns with task behavior, as participants with a larger rise in SCE exhibit better clustering of recalls and superior memory. Finally, two analyses provide evidence that ripples signal context reinstatement. First, ripples during words shown early in the list lead to ripples during presentation of semantically-related words many seconds later in the list and, combined, predict increased recall of these later words. Second, when ripples occur during encoding of a word, that word leads to clustering significantly more often when a ripple also occurs prior to its recall. These findings, in which hippocampal ripples during memory formation predict subsequent ripple-mediated reinstatement during both later list items and retrieval, suggest ripples specifically signal encoding and reinstatement of episodic memories.

During free recall, hippocampal ripples occur just prior to the retrieval of a previously studied item, termed the pre-retrieval effect (PRE) (*4*). The strongest PRE occurs prior to pairs of recalls bearing strong temporal and/or semantic relations, suggesting that hippocampal ripples reflect an item-to-context reinstatement process (*35*). A recent review hypothesizes that sharpwave ripples perform a dual function by mediating both memory formation and retrieval (*2*), as repetition in support of consolidation (*10*) may share mechanisms with reinstatement during retrieval. In light of this hypothesis and the ripple SCE results (**Fig. 3**), we ask if the SCE relates to the PRE. Our final analysis substantiates the hypothesis: recalls with ripples during both the initial word presentation and in the PRE window lead to clustering significantly more than recalls without ripples in both periods (**Fig. 6**). In other words, both the SCE and the PRE appear to reflect a related process, where items bind to context during encoding and subsequently reinstate context from items during retrieval (*35*). Further, considering that participants have prior knowledge of the semantics of the common nouns used in this study and that 46% of recalls lead to clustering (**Fig. 1c**), the SCE also may reflect reinstatement of categorical context (*33*) during word presentation (e.g. sea animals in **Fig. 1b-c**). That participants with larger ripple SCEs show more subsequent clustering of recalls (**Fig. 4b**) supports this interpretation. **Figure 5** also supports categorical reinstatement during encoding, as ripples during a semantic category early in a list *X*_1_*_−_*_2_ predict better recall of words from that same category shown later in the list (*X*_3_*_−_*_4_). This effect does not occur the other way around, as *X*_3_*_−_*_4_ ripples do not increase *X*_1_*_−_*_2_ recalls, suggesting context must reinstate (i.e. during *X*_1_*_−_*_2_) prior to the improvement in encoding (i.e. during *X*_3_*_−_*_4_). Participants recall the most *X*_3_*_−_*_4_ words when ripples occur during both *X*_1_*_−_*_2_ and *X*_3_*_−_*_4_, supporting the idea that context reinstatement during both periods optimizes word encoding.

A more conservative interpretation of the SCE is that it simply reflects engagement of the hippocampal memory system. Tasks with larger memory demands more likely recruit hippocampal involvement. For example, studies of hippocampal amnesics on delayed memory tasks found that deficits only occur if task demand is sufficient (e.g. relatively large set size or retention delays (*36*)). And when MTL amnesics performed a delayed free recall task similar to ours they specifically showed deficits in reinstating context compared to healthy controls, but no difference recalling the most recently-shown items, suggesting deficits occur due to defects specific to the episodic system (*37*). Similarly, single unit recordings support the idea that recruitment of the hippocampus only occurs with sufficient task demands, as hippocampal neurons fail to fire above baseline levels until memory demands are relatively large (*38, 39*). Therefore, when participants engage their hippocampal memory system, whether through increased attention or by forming associations between words from semantic categories, the ripple SCE may reflect the increase in hippocampal activity. Indeed, the SCE increases for participants with higher recall rates (**Fig. 4a**), which suggests that participants successfully recruiting their episodic system show improved memory. And in the case of **Fig. 6**, where ripples mediate episodic encoding and retrieval, in both cases we expect the hippocampus to be engaged as participants learn semantically-associated items during encoding and subsequently recall semantically-associated items during retrieval.

Two previous works have reported a ripple SME in humans. The first suggests that ripples rise only after offset of the initial presentation of face or place pictures that participants subsequently free recall (*5*). We find no evidence for this phenomenon, although unlike our task each picture was shown three addition times, suggesting repetition might influence the initial presentation SME. The second work utilized a cued recall task where participants viewed a face with a written profession underneath, they were asked to say the profession aloud to promote a mental association, and then subsequently recalled the profession when presented the face (*7*). They find a ripple SME from 750-1375 ms after initial presentation of the pair. We believe our finding of a ripple SCE but not a ripple SME accords with this work, as an SCE will manifest as an SME without without having a behavioral contrast to separate episodic from non-episodic retrieval. Their presentation of a face with a profession promotes creation of an associative context that subsequently reinstates when participants see the face during retrieval. Or, considering our more conservative interpretation of the SCE, the creation of the face-profession association promotes hippocampal engagement during encoding. In either case a ripple SCE may be underlying the SME. Finally, we also hypothesized a ripple SME in our OSF registration after the first 35% of participants showed a significant rise in ripples for recalled compared to recalled words (https://osf.io/e98qp) that did not remain after unlocking the full dataset. Our modeling results in **Eq. 4**, where we break down this simpler recalled vs. not recalled contrast into recalled, clustered, and not recalled words, helps us understand why. Words leading to subsequently clustering show a positive coefficient (i.e. have more ripples than not recalled words) while words leading to unclustered recalls show a negative contribution (i.e. have fewer ripples than not recalled words). Combining across all recalled vs. not recalled words for the “overall SME”, the positive contribution from the subsequently clustered recalls appears to outweigh the negative contribution from the remaining recalls for this initial set of participants. But as we increase the sample size to the full dataset, we gain better precision of the standard error estimation, and this “overall SME” fails to remain significant.

We replicate previous work (*7,16,32*) showing high frequency activity (HFA) SMEs throughout the MTL, as each region we test has significantly stronger signal for subsequently recalled than not recalled words. The ubiquity of the HFA SME throughout the MTL is possibly of physiological relevance, as high gamma, which largely overlaps with HFA, is thought to synchronize regions during cognitive tasks (*40*). Surprisingly, and contrary to a hypothesis from our pre-registration (https://osf.io/e98qp), we do not find a significant ripple SME in either of the hippocampal subfields we test **Fig. 2b**. And while ripples during presentation of subsequently recalled words vs. not recalled words peak *∼*0.6 s as shown in the PSTHs for CA1 and CA3/DG, even when we use a narrower 0.4 to 1.1 s window, we still do not find a significant ripple SME in either (*P>*0.30, each FDR-corrected across 4 tests of Eq. 3). These results suggest the algorithms designed to detect ripples in rodents (*41*) achieve a level of specificity that separates ripples from more ubiquitous high frequency signals. What differences in the detector for ripples vs. HFA account for this specificity? Two components are likely to be responsible. First, the ripple detector only considers “candidate” events with power exceeding a high threshold (3 SD). Second, the detector requires these candidate events stay above a lower threshold (2 SD) for a minimum duration (20 ms) to be considered a ripple. Therefore, we speculate that high frequency activity that does not reach sufficiently high powers or arises only transiently accounts for the HFA SME. Future work splitting individual events into ripple vs. HFA groups will be necessary to test these hypotheses.

The present report argues that hippocampal ripples signal the encoding of episodic memories, as the presence of ripples during item encoding predicts the subsequent, ripple-mediated reinstatement of context during retrieval. Considering the specificity in which hippocampal ripples signal this subsequent clustering effect **Fig. 3b**, as opposed to the more ubiquitous HFA subsequent memory effect found throughout MTL **Fig. 2c** and other regions (*16*), future work might take advantage of ripples as a biomarker of episodic memory formation. In particular, considering that classification of brain states that predict memory encoding can be used to time stimulation for the purpose of improving memory (*42*), future work might incorporate ripple detection to specifically target episodic memory formation for use in translational work.

## Acknowledgements

Data were collected as part of the DARPA RAM program (Cooperative Agreement N66001-14-2-4032). This work is supported by NIH Grant R01NS106611 and US Army Medical Research and Development Command Medical Technology Enterprise Consortium Grant MTEC-20-06-MOM-013. The views, opinions, and/or findings contained in this material are those of the authors and should not be interpreted as representing the official views or policies of the Department of Defense or the US Government. We thank the Kahana Lab and the Joshua Jacobs Lab for providing valuable feedback on this work.

